# Single cell analysis of human CD8^+^ T cells reveals CD45RC^low/-^ TNFR2^+^CD29^low^CD8^+^ Tregs with superior activity

**DOI:** 10.1101/2023.12.13.571475

**Authors:** Céline Sérazin, Léa Flippe, Mathias Streitz, Désirée-Jacqueline Wendering, Stephan Schlickeiser, Frederik Heinrich, Pawel Durek, Gabriela Guerra, Katrin Lehmann, Mir-Farzin Mashreghi, Harald Wajant, Hans Dieter Volk, Ignacio Anegon, Laurent David, Séverine Bézie, Carole Guillonneau

**Author notes:** **Corresponding author:** Dr. Carole Guillonneau, INSERM UMR1064 – Center for Research in Transplantation and Immunology-ITUN, 30 Bd Jean Monnet, 44093, Nantes Cedex 01, France.

## Abstract

Although described in the 70’s, CD8^+^ regulatory T cells (Tregs) remain incompletely understood and to date, although several markers are used to define them, they remain poorly defined. The identification of reliable and consistent markers, as it was done for CD4^+^ Tregs, remains an urgent task and a challenge to advance our understanding. Herein, we analyzed total CD8^+^ T cells using single cell CITEseq and VDJ T cell receptor sequencing utilizing markers used previously to identify Tregs, in particular CD45RC described by our team and others to divide pro-inflammatory (CD45RC^high^) and pro-regulatory (CD45RC^low/-^) CD8^+^ T cells in rat, mice and human. 7000 freshly isolated, non-stimulated CD8^+^ T lymphocytes of four healthy volunteers were analyzed. Combining at a single cell level transcriptome and protein expression data led for the first time to the characterization and definition of three subsets of regulatory CD8^+^ T cells. Further *in vitro* functional analysis based on three markers highlighted the superior suppressive activity of the CD8^+^CD45RC^low/-^TNFR2^+^CD29^low^ Tregs subset.

To our knowledge, this is the largest characterization of human CD8^+^ Tregs to date. This data resource will help improve our understanding of CD8^+^ T cells heterogeneity and will help to translate CD8^+^ Tregs to the clinic.

## INTRODUCTION

Regulatory T cells are key components of the immune system involved in the maintenance of tolerance and acting to suppress immune responses in many pathological situations. CD4^+^ Tregs are the most described immunosuppressive cells since the discovery in 1995 of CD25 as a cell membrane marker for their identification correlating with the discovery of a Treg specific transcription factor forkhead box protein P3 (FOXP3)^1^. Since then, numerous efforts have been made for the comprehension of these cells critical in the regulation of the balance of the immune system, including the characterization of their origin and their transcriptional landscape in various compartments. At the opposite, very few efforts have been done for a better understanding of CD8^+^ Tregs (or CD8^+^ T suppressor cells as they were called in 1972 ^2^) despite having been described by many groups in multiple pathophysiological situations in different species including humans ^3–5^. The role for FOXP3 in CD8^+^ T cells has been poorly described and is still controversial, similarly the existence of a CD8^+^ Tregs lineage specific transcription factor is still debatable, which together with the absence of consensual and specific surface markers, led to the general disregard of CD8^+^ Tregs.

The rapid development in the last years of technologies and methods to gain detailed information on the transcriptome and proteome at a single cell level has allowed the discovery of rare cell populations, as well as their lineage specifications. However, to date, most studies on regulatory cells have focused on CD4^+^ Tregs, and there has been no true attempt to reveal CD8^+^ Treg subsets and understand the heterogeneity within them and relationship between CD8^+^ T cell subsets. Indeed, as for other cell populations, one could expect that CD8^+^ Tregs exist in different identities related to distinct functional characteristics and differentiation pathways.

So far, our team ^6–10^ and others ^11–13^ have demonstrated that CD8^+^ Tregs identified by low and/or negative expression of the marker CD45RC, one of the isoforms of the CD45 molecule, show potent suppressive activity *in vitro* and *in vivo,* while mouse, rat and human cells expressing high levels of CD45RC do not. CD45RC has been revealed as a critical marker to distinguish within both CD4^+^ ^13^ and CD8^+^ T cells ^11^ the pro-inflammatory cells (CD45RC^high^) from pro-regulatory cells (CD45RC^low/-^). Based on these observations, since more than 15 years, our team isolates CD8^+^ Tregs based on the following phenotype: CD3^+^CD56^-^ CD8^+^CD45RC^low/-^ since no better marker could be evidenced. We have demonstrated the suppressive function of these cells freshly isolated and/or expanded for up to 21 days *ex vivo* and their potential to prevent transplantation rejection, GvHD and multiple sclerosis ^3,6–8,14–17^. We demonstrated that expanded CD8^+^CD45RC^low/-^ Tregs show an increased suppressive activity compared to the fresh one and differences in their phenotype, notably the overexpression of FOXP3 ^7^. According to phenotype analysis by flow cytometry, CD8^+^CD45RC^low/-^ Tregs is a heterogeneous population based on surface markers in peripheral blood, and this heterogeneity is decreasing with polyclonal ex vivo expansion to reach to some extent a homogenous and clinically applicable population. It suggests that either a fraction of the circulating CD8^+^CD45RC^low/-^ population is tolerogenic, comes from the thymus and is preferentially expanded, or that the induction of tolerance is peripheral and due to a combination of cells forming an "immunological niche" and allowing for the generation of induced CD8^+^CD45RC^low/-^ Tregs.

Systematic evaluation in fresh CD8^+^CD45RC^low/-^ Tregs of proposed markers by flow cytometry, including CD28, CD122 and else, was proven unsuccessful to enrich the tolerogenic activity ^7^. Thus, a deep characterization of fresh CD8^+^CD45RC^low/-^ T cells is needed for delineating their heterogeneity and to clearly define a strong and accepted consensus and solve the ongoing debate on the phenotype of CD8^+^ Tregs mostly with the finding of new highly specific markers.

Here, we employed single cell RNA-seq coupled to CITEseq including VDJ sequencing in order to precisely delineate CD8^+^ Tregs in human peripheral blood. With this approach we could nail down three subsets of Tregs, which allowed us to gain unprecedented knowledge of those key immunological players. Finally, we identified surface markers predicting higher Treg activity and validated two of them in functional assays. Altogether our study will greatly improve our ability to understand the role of CD8^+^ Tregs in immunological response.

## RESULTS

### Single cell RNA-seq coupled to CITE-seq mapped CD8^+^ T cells and resolved subsets distribution

In order to prevent individual-biased results, we labelled with barcoded antibodies (HTOs) and pooled freshly isolated CD8^+^ T cells (CD4^−^ CD3^+^) from the blood of 4 healthy volunteers (**Suppl. Figure 1A**). The cells were analyzed by single-cell RNA-seq coupled with 30 CITE-seq antibodies and, to account for CD8^+^ regulatory T cells, we included several markers related to Tregs including CD45RC, since isoforms of CD45 could not be distinguished by transcriptomic analysis (**Figure 1A**). CD8^+^ T cell purity was checked at transcriptomic and proteomic level and we excluded NKT cells (CD3^+^CD56^+^CD16^+^) of our analysis (**Suppl. Figure 1B-C**).

**Figure 1.**
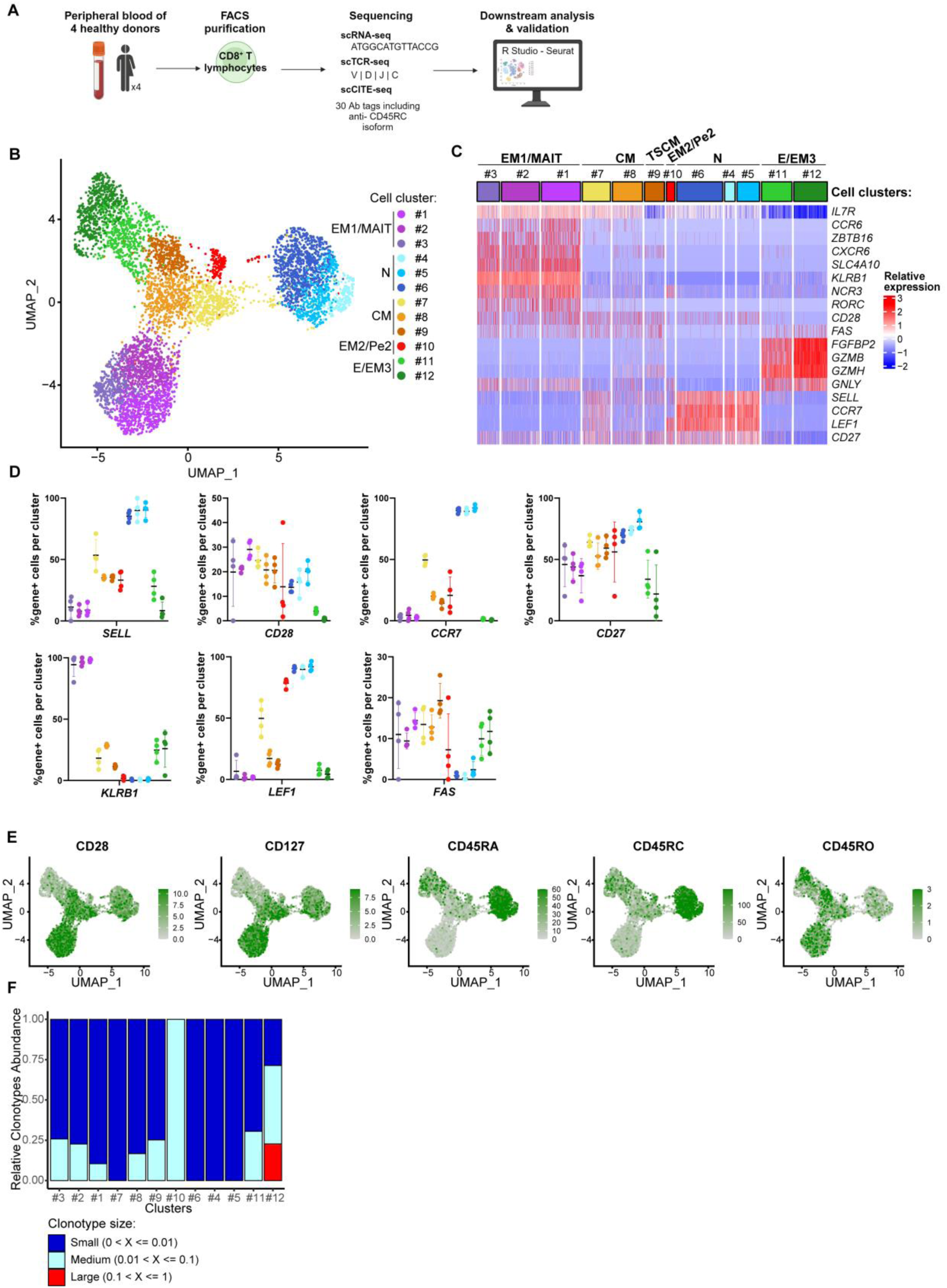
Single cell RNA-seq coupled to CITE-seq mapping of CD8^+^ T cells. **(A)** Schematic representation of 5’ single cell RNA-sequencing workflow (created with Biorender). **(B)** Identification of CD8^+^ T cell subpopulations with the definition of twelve cell clusters on the Uniform Manifold Approximation and Projection (UMAP) based on genes expression (resolution = 1). Naive (N), central memory (CM), effector memory (EM) type 1 (EM1)/ mucosal invariant T cell (MAIT), type 2 (EM2), type 3 (EM3) and pE2 (TEMRA) CD8^+^ T cells have been identified. Each dot corresponds to a single cell. **(C)** Heatmap of selected genes for defining cell cluster’s identity. Columns correspond to single cells grouped by cell clusters and rows correspond to genes. Expression values were scaled per gene. Blue color represents lower expressed genes and red color represents higher expressed genes. **(D)** Frequencies of cells expressing each gene in CD8^+^ T cells clusters. Each dot corresponds to an individual. Each color corresponds to a cluster. **(E)** Distribution of CD28, CD127, CD45RA, CD45RC and CD45RO protein detection across clusters identifies CD45RC^neg^, CD45RC^low^ and CD45RC^high^ clusters. **(F)** Distribution of clonotype across cell clusters. In red are the clonotypes found in from 10 to 100% cells in the cell cluster, in light blue from 1% to 10% cells of the cell cluster and in dark blue from 0 to 1% cells of the cell cluster.

RNA-seq data of a total of 7517 single cells evenly distributed among healthy individuals meeting the inclusion criteria (**Suppl. Figure 1D-F**) were integrated in Seurat, log normalized and visualized by Uniform Manifold Approximation and Projection (UMAP). We confirmed that there was no major visible bias (number of reads and mitochondrial genes content) and that reads passed quality check (see mat & met). To browse the data, we generated a first clustering that defined 12 distinctive cell clusters (**Figure 1B**).

Expression of genes (*SELL*, *CCR7, CD27* and *LEF1*) and surface markers (CD45RA and CD28) identified naive T cells (cell clusters 4, 5 and 6) (**Figure 1B-E**) while effector memory 1 (EM1) stage could be identified by the higher level of *ZBTB16*, *RORC*, *SLC4A10*, *KLRB1, NCR3, CD28, CD27, IL7R* and low/no expression of *CCR7,* with a CD45RA^-^ CD127^+^ CD28^+^ phenotype (**Figure 1B-E**) (cell clusters 1, 2 and 3). We observed that cell clusters 1, 2 and 3 showed a MAIT cell profile with expression of *TRAV1-2*/*TRAJ33* TCR chains (Suppl. **Figure 1G**), *KLRB1* (encoding CD161), *SLC4A10*, *CCR6* and *CXCR6* genes ^18^ and a memory EM1 phenotype (i.e. CD95^hi^CD62L^lo^CD45RO^+^CD45RA^lo^CD27^+^ ^19^). The remaining cell clusters could not be identified using only T cell known surface markers, therefore we also considered expression of key developmental genes. This allowed us to subcategorize cell clusters #11 and #12 as E/EM3 T cells (on the higher expression of *FGFBP2*, *GZMB*, *GZMH*, low expression of *CD27* and no expression of *CCR7* and CD45RA^low/-^CD28^low/-^ phenotype), cell clusters 7-9 and 10 as central memory (CM) stage (positive expression of *CCR7* and *CD27* and CD45RA^-^ CD28^+^ phenotype) and EM2/pE2 stage (positive expression of *CD27* and *CCR7*, and a CD45RA^low/-^CD28^-^ phenotype) respectively. The CM cluster could be further subcategorized by the expression of CD95 (*FAS*) to allow identification of stem cell-like memory T cells (TSCM) in cluster 9.

We then further analyzed on the different clusters identified the different levels of expression of CD45RC i.e. high, low or negative based on CITE-seq antibody expression (**Figure 1E and Suppl. Figure 2A-G**). We observed that the CD45RC marker allowed a clear distinction between cell clusters 4, 5 and 6 displaying a high expression of CD45RC and the remaining cell clusters displaying a low or negative expression (**Suppl. Figure 2A-D**). The proportion of CD45RC cells expressing high, low or negative levels was equally distributed among donors (**Suppl. Figure 2E**). In addition, analysis of genes differentially expressed between cells expressing high, low or negative levels of CD45RC evidenced that cells expressing low or negative levels of CD45RC are close in transcriptomic profile while CD8^+^CD45RC^high^ T cells are clearly distinct (**Suppl. Figure 2F-K**). Since we and others previously demonstrated that CD8^+^CD45RC^high^ T cells do not exert suppressive activity, it suggest that the pro-regulatory cells are contained within the remaining cell clusters i.e. 1-3, and 7-12 ^7,8,14,17^.

Finally, the relative abundance of TCR clonotypes per cell cluster was analyzed based on the frequency of clonotypes ranging from small (<0.01), medium (between 0.01 and 0.1) and large (>0.1). Most cell clusters demonstrated small clonotype frequency, however we observed that cell clusters 10, 11 and 12 demonstrated more frequent clonotypes (**Figure 1F**). These findings suggest that these cell clusters (i.e. 10, 11 and 12) are dominated by some expanded clonotypes compared to the other cell clusters, correlating with the memory status of the cells defined within (**Suppl. Table 1**). Except from these clusters, at this level of analysis, we did not find specific TCR signatures for the other clusters.

As suppressive assays on subpopulations of CD161^high/int/-^ demonstrated absence of suppressive properties for MAIT cells (cell clusters 1,2 and 3) (**Data not shown**), they were thus excluded from further analysis of the transcriptomic data.

### Identification of 3 populations within CD45RC^low/-^CD8^+^ T cells with regulatory signatures

To further characterize the heterogeneity within CD8^+^ T cells, we projected the cells by UMAP, excluding the MAIT cells defined previously, allowing now a definition of 10 new cell clusters (**Figure 2A**). We observed that the distribution according to T cell development stage was similar to what we previously observed (**Suppl. Figure 3A**). A total of 2314 genes were differentially expressed between cell clusters allowing better definition of each cell clusters with more than 80 and up to 540 genes differentially expressed genes per cell cluster (**Figure 2B**). Analysis using the CD45RC marker showed a total of 1179 genes differentially expressed between CD8^+^CD45RC^high^ (clusters 0, 5 and 7) and CD8^+^CD45RC^low/-^ (clusters 1, 2, 3, 4, 6, 8 and 9) T cells (**Figure 2C**), that were evenly distributed among healthy individuals (**Suppl. Figure 3B**). Volcano plot analysis comparing CD8^+^CD45RC^high^ and CD8^+^CD45RC^low/-^ T cells (**Figure 2C**) together with heatmap analysis of classical T regulatory cells-associated genes (**Figure 3A-B**) highlighted some signature genes and demonstrated the lower expression of regulatory genes in the CD8^+^CD45RC^high^ T cells. Indeed, the gene signature of CD8^+^CD45RC^high^ T cells was associated with significant higher expression of membrane proteins such as *MAL*, *LRRN3* and *CCR7* or transcription factors such as *LEF1* (**Figure 2C**). Analysis of total CD8^+^CD45RC^low/-^ T cells highlighted significant higher expression of some genes encoding membrane proteins (*CD99*, *KLRG1*, *PLEK*, *KLRD1* (i.e. CD94), *IL10RA*, *IL2RB*, *ITGB1* and *TNFRSF1B* (i.e. TNFR2)) and secreted proteins (*CCL5*, *CST7*, *GZMA* and *TGFB1*) or transcription factors such as *IKZF2* (i.e. HELIOS) compared to CD8^+^CD45RC^high^ T cells (**Figure 2C**). Expression of *FOXP3* in contrast was not differentially expressed between cell clusters and remained rarely detected (**Suppl. Figure 3C**), correlating with studies demonstrating the low expression of FOXP3 by unstimulated fresh CD8^+^ T cells ^7^. Within CD8^+^CD45RC^low/-^ T cells some cell clusters were thus of particular interest, namely cell cluster 8 and 9 which represented small cell clusters in size with 146 and 44 cells, respectively showing a higher expression of *IKZF2* (HELIOS) and cell clusters 1/2 showing a higher expression of *TNFRSF1B* (TNFR2) (cluster 2 in particular) and a lower expression of *IL7R* (CD127) and *CD28* **(Figure 2D)**. Finally, we observed expression of *FGL2* in cell cluster 2, a gene associated to CD4^+^ Treg suppressive function and that we have also previously described as involved in CD8^+^ Treg suppressive function ^20^.

**Figure 2:**
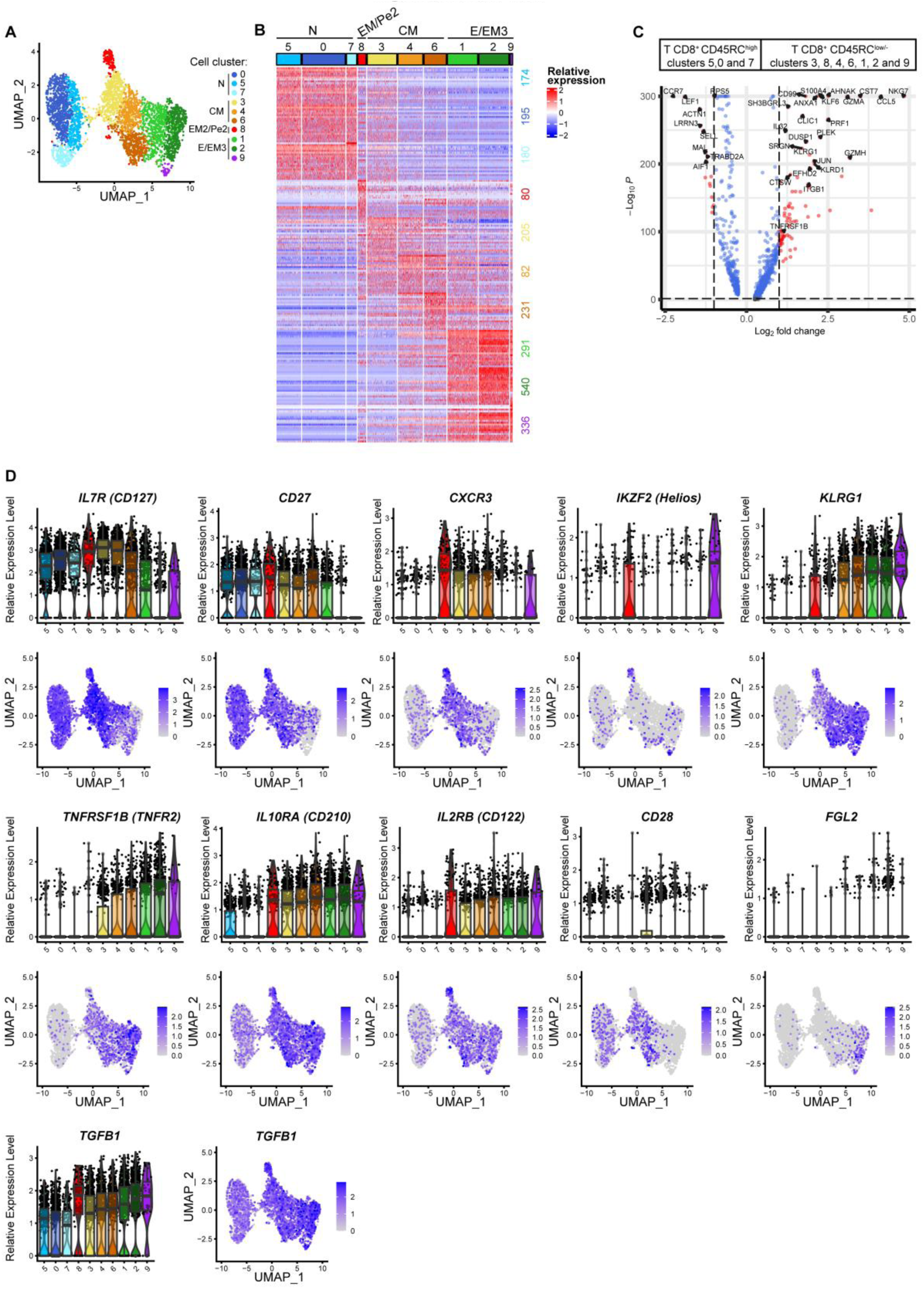
Characterization of CD8^+^ regulatory cell clusters. **(A)** Definition of 10 cell clusters in fresh CD8^+^ T cells after exclusion of EM1/MAIT cells cluster on the Uniform Manifold Approximation and Projection (UMAP) based on gene expression (resolution = 0.8). Each dot corresponds to a single cell and each color to one cell cluster. **(B)** Heatmap of the top 20 differentially expressed genes (min.pct = 0.25, logfc.threshold = 0.2) per cell cluster. Expression values were scaled per gene. Blue color represents genes with lower expression and red color represents genes with higher expression. The total number of genes characterized in each cell cluster is provided on the right, colors correspond to cell cluster. **(C)** Volcano plot highlighting significant differences (min.pct = 0.25, logfc.threshold = 0.2) in gene expression between CD45RC^high^ (cluster 5, 0 and 7, left) vs CD45RC^low/-^ (clusters 3, 8, 4, 6, 1, 2, 9, right) CD8^+^ T cells. The thresholds have been set to p value adjusted <0.05 and fold change >1.5. Red dot corresponds to genes that have exceeded both thresholds and blue dot corresponds to genes with a significant p-value adjusted. Labels were added to *TNFRSF1B* and the top most differentially expressed genes. **(D)** Feature plots and violin plots of genes related to Tregs. For feature plots, gene expressions are scaled from grey to blue and each dot corresponds to a single cell. For violin plots, gene expression is represented in each cell clusters. Colors correspond to cell cluster and each dot corresponds to a single cell.

**Figure 3:**
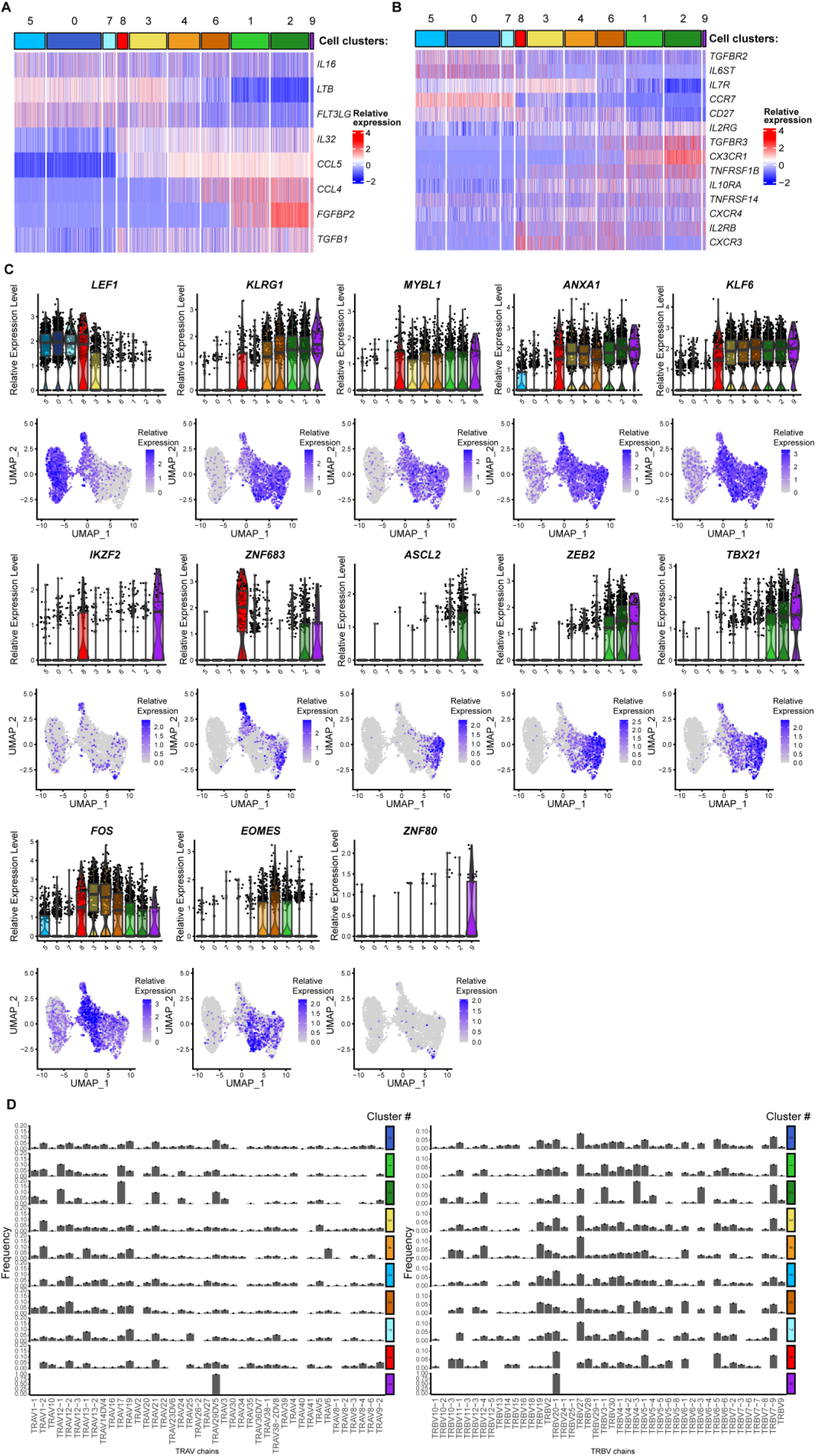
Characterization of CD8^+^ regulatory cell clusters. **(A)** Heatmap of genes encoding for cytokines receptor that are differentially expressed between cell clusters (min.pct = 0.25, logfc.threshold = 0.2). Columns correspond to cell clusters and rows correspond to genes. Expression values were scaled per gene. Blue color represents less expressed genes and red color represents more expressed genes. **(B)** Heatmap of genes encoding for cytokines that are differentially expressed between cell clusters (min.pct = 0.25, logfc.threshold = 0.2). Columns correspond to cell clusters and rows correspond to genes. Expression values were scaled per gene. Blue color represents less expressed genes and red color represents more expressed genes. **(C)** Feature plots and violin plots of transcription factors. **(D)** Frequency of the relative usage of variable domain of (left) TRA or (right) TRB chains in each cell clusters.

Further analysis of cytokine/chemokine receptor expression (**Figure 3A**) highlighted higher expression of *TNFRSF14* (HVEM), *CXCR3, TGFBR3* in addition to *TNFRSF1B* (TNFR2) by cell cluster 1, 2 and 9. *IL2RB* and *CXCR3* were more expressed by cell cluster 8. Cell cluster 0, 5 and 7 showed higher expression of *CCR7*, *CD27*, *IL6ST* and *TGFBR2*.

Analysis of cytokines/chemokines signature genes (**Figure 3B and Suppl. Figure 3D**) demonstrated a similar pattern between cell clusters 1, 2 and 9 expressing similar cytokines (*FGFBP2*, *CCL4*, *CCL5*, and *TGFB1*). Cells of cluster 8 expressed higher levels of *IL32*, *TGFB1*, *LTB* and *IL16*, while cells of clusters 0, 5 and 7 expressed higher levels of *FLT3LG*, *LTB* and *IL16* than cells of the other clusters. Surprisingly, we observed no specific association between expression of senescence/activation or exhaustion associated markers and any of the cell clusters (**Suppl. Figure 3E**).

Finally, we investigated the expression of lineage-defining transcription factors for each cell cluster and projected on a heatmap and feature plot the genes differentially expressed between cell clusters (**Figure 3C and Suppl. Figure 3G**). Interestingly, we were able to identify specific signatures for each cell cluster. We observed a common signature of transcription factors for all CD8^+^CD45RC^high^ T cells (clusters 0, 5, 7), in particular higher expression of *LEF1* essential for repressing CD4^+^ lineage-associated genes including *CD4*, *FOXP3* and *RORC* in CD8^+^ T cells ^21^. In contrast, we found that *KLRG1*, *MYBL1*, *ANXA1* and *KLF6* were associated with total CD8^+^CD45RC^low/-^ T cells. Cell clusters 8 and 9 show a common expression of *IKZF2*, a transcription factor already associated to regulatory function of both CD4^+^ and CD8^+^ Tregs ^5^. Cell cluster 8 also share with cell clusters 2 and 9 the expression of *ZNF683*, a tissue resident T-cell transcription regulator. We observed that cell cluster 1/2 has a similar signature although the one of cell cluster 2 was more pronounced with for example expression of *ASCL2*, *ZEB2* or *TBX21*, transcription factors associated to activated T cell and polarized Tregs ^22^. We observed the higher expression of *FOS* in cell cluster 3/4 and of *EOMES* in cell cluster 6. Finally, cell cluster 9 also displayed a unique transcriptional signature with expression of *ZNF80* in addition to the previously mentioned expression of *IKZF2*.

Since natural Tregs selected in the thymus have been shown (at least for CD4^+^ Tregs) to express TCRs that are distinct to some extent from the TCR repertoire of effector T cells, we sought to further determine the TCR alpha and beta chain preferential usage of the different cell clusters and sharing in between them. For this, we assessed the diversity of TCRs and performed a chord plot analysis of the TCR clonotypes (both alpha and beta chains) (**Figure 3D and Suppl. Figure 3F**). Cell cluster analysis per individuals showed a vast and diverse repertoire for all 4 individuals **(Figure supp 3F)**. We also observed that most cell clusters demonstrated some sharing of the clonotypes, we observed the most important sharing of the clonotypes in cell clusters 1/2 for all the 4 donors, 8 for 2 out of 4 donors, and 9 for 1 donor and less sharing for cell clusters 0, 5 and 3. Usage of TCR alpha and beta chains demonstrates a clonal expansion for cell cluster 9 whereas for other cell cluster no bias was found **(Figure 3D)**.

Given these results, we speculated that based on unsupervised clustering of single cell RNA-seq data, several distinct human CD8^+^ Treg subsets can bet defined at steady state and could correspond to cell clusters 1/2, 8 and 9 and exhibit distinct status of maturation and possibly function. Although cell cluster 9 share similarities with clusters 1 and 2, this cluster is specific of only one donor and display a unique TCR repertoire, thus it was considered as distinct from clusters 1 and 2. We excluded the possibility that mature and functional CD8^+^ Treg subsets might exist within the remaining cell clusters.

### Characterization of CD8^+^ T regulatory cell clusters highlights potential markers including *TNFRSF1B* and *ITGB1*

To better comprehend the specific features of the cell clusters contained within the CD8^+^CD45RC^low/-^ T cells and in particular for cell clusters with demonstrated potential interest as human CD8^+^ Treg subsets i.e. cell clusters 1/2, 8 and 9, we excluded the CD8^+^CD45RC^high^ T cells and projected the CD8^+^CD45RC^low/-^ T cells by principal component analysis, visualized by UMAP (**Figure 4A and Suppl. Figure 4A**) and compared by volcano plot the transcriptomes (**Figure 4B-E**) of the cell clusters of interest vs the remaining cell clusters. The analysis allowed identification of 7 distinctive cell clusters on the UMAP (**Figure 4A**). To facilitate the comprehension, we attributed to the new cell clusters the same numbers and colors as described in **Figure 2A**. We then projected on a volcano plot the genes differentially expressed between cell cluster 1/2 and the remaining cell clusters (**Figure 4B**), cell cluster 8 and the remaining cell clusters (**Figure 4C**) and cell cluster 9 and the remaining cell clusters (**Figure 4D**). We found overall that each of these specific cell clusters (i.e. 1/2, 8 and 9) showed a specific gene signature possibly associated with Tregs using different mechanisms of suppression or different suppressive capacity, and suggesting that Tregs may be presented in each of these cell clusters. Further analysis of genes differentially expressed in these cell clusters revealed several interesting genes already associated with Treg and tolerance for cell cluster 1/2 such as *ITGB1*, *TNFRSF1B*, *GPR56* and *CX3CR1* (**Figure 4B**). In cell cluster 8, we found the interesting expression of *FUT7*, a gene mediating synthesis of CD15s, a molecule expressed in eTreg and identifying the most suppressive Tregs ^23^. Analysis of cell cluster 9 demonstrated the specific expression of *KIR2DL3* (**Figure 4D and Suppl. Figure 4B**), suggesting altogether with *IKZF2* expression, that this signature is related to human CD8^+^ Tregs described in anti-viral immune responses based on recent work published ^4^ and that the cell cluster 9 could correspond to a CD8^+^ Treg subset involved to suppress pathogenic T cells in infectious diseases.

**Figure 4:**
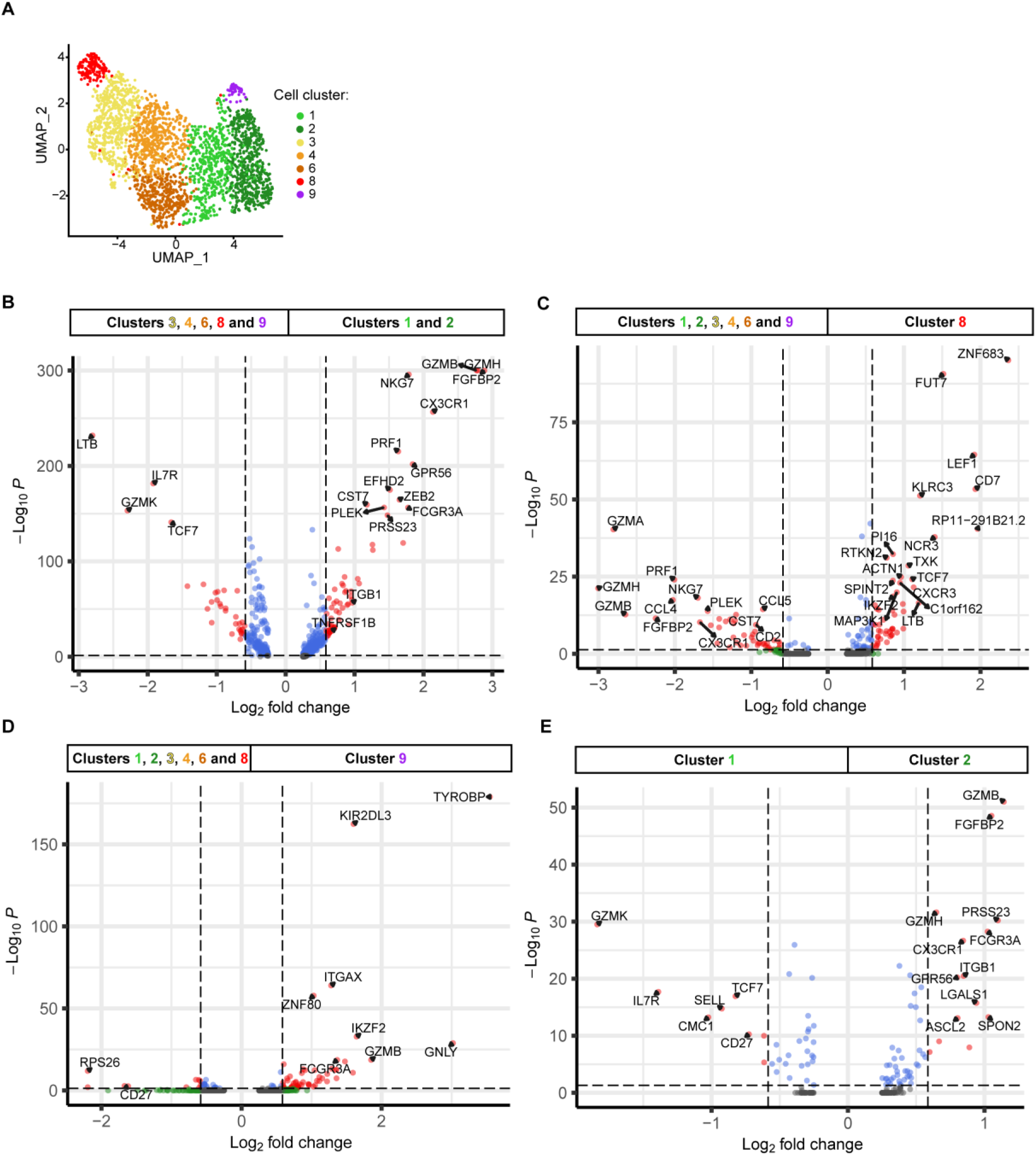
Identification of markers specific of CD8^+^ T regulatory cells clusters. **(A)** Definition of 7 cell clusters in fresh non MAIT CD8^+^CD45RC^low/-^ T cells on the Uniform Manifold Approximation and Projection (UMAP) based on gene expression (resolution = 0.5). Each dot corresponds to a single cell. Cell cluster numbers and colors are conserved from Figure 2A. **(B-E)** Volcano plot showing differentially expressed genes between cell cluster 1 and 2 vs all other cell clusters **(B)**, between cell cluster 8 vs all other cell clusters **(C)**, between cell cluster 9 vs all other cell clusters **(D)**, between cell cluster 1 vs cell cluster 2 **(E)**. The thresholds have been set to p value adjusted <0.05 and fold change >1.5. Red dot corresponds to genes that have exceeded both threshold, and blue dot correspond to genes that have a significant p-value adjusted but not fold change. *TNFRSF1B* and *ITGB1* genes and the top most differentially expressed genes were labelled.

To identify genes deciphering cell cluster 1 from cell cluster 2, we analyzed their transcriptomic differences using a volcano plot (**Figure 4E**). We found that cells from cluster 2 exert a differentiated and effector profile with expression of genes such as *FGFBP2*, *CX3CR1* and *FCGR3A* ^24^, as well as genes previously mentioned as such as GPR56 ^25^ and ITGB1^26^. In cell cluster 1 we found *GZMK*, *IL7R*, *SELL*, *CD27* and *TCF7* to be differentially expressed corresponding rather to a central memory phenotype. No specific Treg-related genes could be identified in cell cluster 1 vs cell cluster 2.

Altogether this analysis confirms that several potential distinct CD8^+^ Treg subsets exhibiting different transcriptomic signatures might exist in human correlating with description of CD4^+^ Treg cells.

### TNFR2 protein expression defines CD8^+^ Tregs and contributes to functional activity

To further get insight into the suppressive potential of the CD45RC^low/-^CD8^+^ Tregs characterized as cell cluster 2, we selected TNFR2^+^ cells whose protein expression using an anti-TNFR2 CITE-seq antibody projected on the UMAP plot (**Figure 5A and Suppl. Figure 5A-B**) confirmed the gene expression observed in the scRNAseq (**Figure 2-4**). We also analyzed the transcriptomic differences of TNFR2^+^ vs TNFR2^-^ CD8^+^CD45RC^low/-^ T cells using the CITEseq anti-TNFR2 mAb in the scRNAseq dataset using a volcano plot (**Figure 5B**). The analysis confirmed that *TNFRSF1B*, the gene coding for TNFR2, was among the top most differentially expressed genes together with other genes associated with T cell proliferation, migration and activation (**Figure 5B-C**), confirming a correlation with cell cluster 2. Our analysis also identified other genes as positively correlating with *TNFRSF1B*/TNFR2 such as *GPR56*, *CX3CR1*, *S1PR5* or negatively correlating such as *CXCR3, GPR183* or *CD28*, correlating with description of CD28^low/-^ CD8^+^ Tregs ^27,28^ (**Figure 5D).**

**Figure 5:**
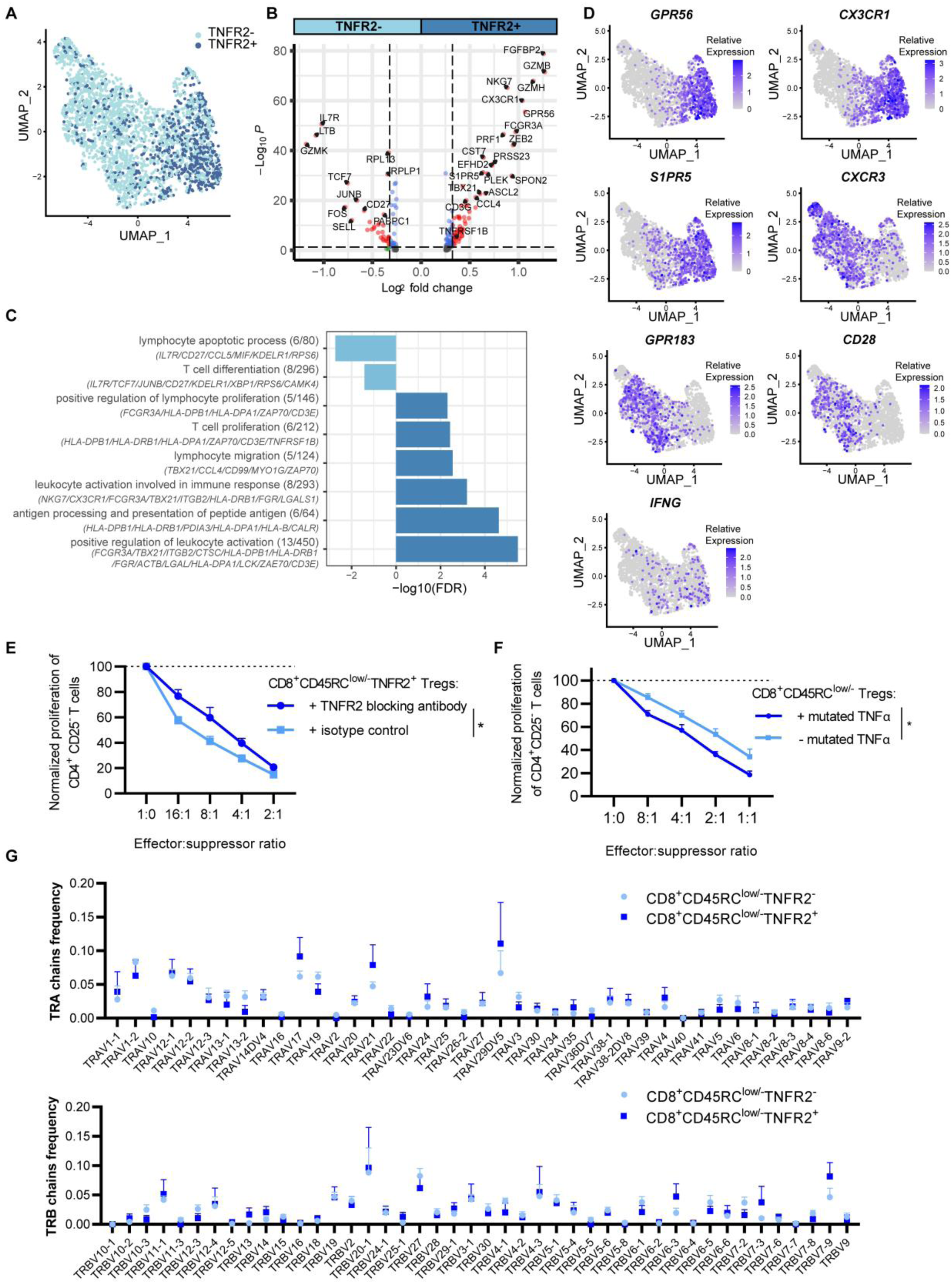
Implication of TNFR2 in suppressive function of CD8^+^CD45RC^low/-^ Tregs. **(A)** Definition of TNFR2^+^ (dark blue) and TNFR2^-^ (light blue) CD8^+^CD45RC^low/-^ T cells with anti-TNFR2 CITE-seq antibody on the UMAP plot. Each dot corresponds to a single cell. **(B)** Volcano plot showing differentially expressed genes between cell cluster TNFR2^+^ and TNFR2^-^ CD8^+^CD45RC^low/-^ T cells. The thresholds have been set to p-value adjusted <0.05 and fold change >1.5. Red dot corresponds to genes that have exceeded both threshold, and blue dot correspond to genes that have a significant p-value adjusted but not fold change. *TNFRSF1B* gene and the top most differentially expressed genes were labelled. **(C)** Normalized enrichment score/FDR of biological pathways upregulated or downregulated in TNFR2^+^ CD8^+^CD45RC^low/-^ T cells compared with TNFR2^-^ CD8^+^CD45RC^low/-^ T cells. **(D)** Feature plot of genes encoding for membrane proteins that correlates or anti-correlates with TNFR2 expression in CD8^+^CD45RC^low/-^ Tregs. Gene expressions are scaled from grey to blue. Each dot corresponds to a single cell. **(E)** The percentage of proliferation of responder T cells in presence of CD8^+^ CD45RC^low/-^ TNFR2^+^ T cells in presence of 10 µg/mL blocking anti-TNFR2 antibody or isotype control in a range of Teff:Treg ratios was normalized to the proliferation of responder T cells in absence of Tregs. Groups were compared using two-way row matched Anova test. *p value > 0.05. Results are expressed as mean ± SEM (n= 4). **(F)** The percentage of proliferation of Teff cells in presence of CD8^+^CD45RC^low/-^ T cells stimulated overnight or not with 100 ng/ml of TNC-scTNF(143N/145R), in culture media during the 5 days of culture in a range of Teff:Treg ratios was normalized to the proliferation of responder T cells in absence of Tregs. Groups were compared using Anova test. *p value > 0.05. Results are expressed as mean ± SEM (n= 5). **(G)** Relative frequency of variable domain of TRA (upper) and TRB (lower) chains in CD8^+^CD45RC^low/-^TNFR2^+^ (dark blue) and CD8^+^CD45RC^low/-^TNFR2^-^ (light blue). Groups were compared using two-way Anova test. Results are expressed as mean ± SEM (n= 4).

To assess the role of TNFR2 as a functional marker and given the described functional relevance of TNFR2 for CD4^+^ Tregs ^29^, we tested the suppressive function of TNFR2^+^CD45RC^low/-^ CD8^+^ T cells in presence of blocking anti-TNFR2 antibody or isotype control antibody (**Suppl. Figure 5C**). The addition of blocking anti-TNFR2 antibody significantly decreased the suppressive activity of TNFR2^+^CD8^+^CD45RC^low/-^ Tregs (**Figure 5E**). This result suggests that TNFR2 signalling is important for the suppressive activity of TNFR2^+^ Tregs. However, as shown previously ^7^, other mechanisms might be of importance, too as inhibition of TNFR2 signals could not fully restore the proliferation of conventional T cells.

We then stimulated the TNFR2 with TNC-scTNF(143N/145R), a nonavalent human TNFR2-specific agonist composed of the tenascin-C trimerization domain genetically fused to a single-chain encoded cassette comprising three human TNF protomers with mutations preventing TNFR1 binding ^30^ (**Figure 5F**). We observed that addition of STAR2 in the suppressive assay led to a significant increase in the suppressive property of total CD8^+^CD45RC^low/-^ T cells on conventional CD4^+^CD25^-^ T cells proliferation, confirming the relevance of TNFR2 for the suppressive activity.

Further analysis of the alpha and beta chain usage of TNFR2^+^CD45RC^low/-^ CD8^+^ T cells vs TNFR2^-^CD45RC^low/-^ CD8^+^ T cells did not reveal a significant bias toward a particular alpha and beta chain, although we observed an increased usage of TRAV17, 21 and 29 and TRBV6-3 and 7-9 (**Figure 5G**). Similarly, analysis of CDR3α and CDR3β amino acid (aa) length, did not reveal any particular bias toward a specific length for TNFR2^+^CD45RC^low/-^ CD8^+^ T cells vs TNFR2^-^CD45RC^low/-^ CD8^+^ T cells (**Suppl. Figure 5D**).

### TNFR2^+^CD29^low^CD8^+^CD45RC^low/-^ T cells are highly suppressive Tregs

Finally, to refine further the phenotype of cells from cell cluster 2 which includes the most of potentially regulatory cells, we considered genes encoding for membrane proteins highlighted by the single cell transcriptomic analysis. We thus selected *ITGB1* (CD29) in addition to *TNFRSF1B* (TNFR2) (**Figure 6A**). At proteomic level, TNFR2 and CD29 surface expression allowed identification of four subsets in peripheral blood CD8^+^CD45RC^low/-^ T cells (**Figure 6B**). Interestingly, we found significatively more CD29^high^TNFR2^-^ cells and less CD29^low^TNFR2^-^ cells in CD8^+^CD45RC^low/-^ T compared to CD8^+^CD45RC^high^ T cells (**Figure 6B**). Although FOXP3 was not specific for cell cluster 2 in the single cell RNA-seq dataset, we further analysed a potential correlation at the protein level as shown in the literature for CD4^+^ Tregs ^31^, and observed a positive correlation of FOXP3 with TNFR2 expression but not CD29 (**Figure 6C**). We then sorted the four subsets on TNFR2^+^ or ^-^ combined with CD29^low^ or ^high^ expression from peripheral blood of healthy subjects and tested their suppressive functions on allogenic CD4^+^ effector T cells stimulated with irradiated allogeneic antigen presenting cells (APC). Interestingly, we found that the CD8^+^CD45RC^low/-^ TNFR2^+^ T cell subsets (both CD29^high^ and CD29^low^) were significantly more suppressive than the total CD8^+^CD45RC^low/-^ T cell population (**Figure 6D**), pointing to a role for TNFR2 in CD8^+^ Treg development or function. We observed a trend for even a stronger suppressive activity for the TNFR2^+^CD29^low^ Treg compared to the TNFR2^+^CD29^high^ subset.

**Figure 6:**
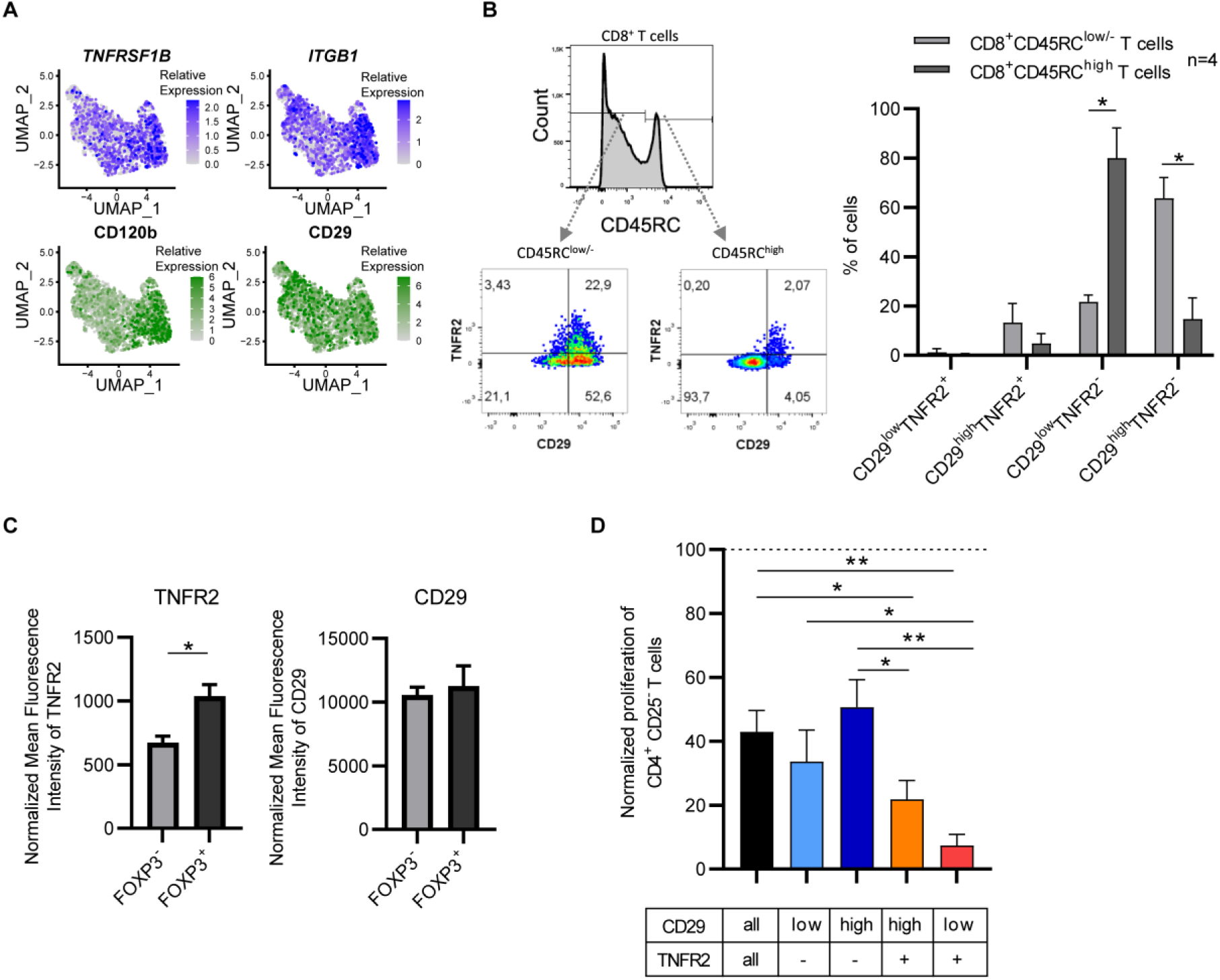
Isolation of CD8^+^CD45RC^low/-^ T cells on TNFR2^+^CD29^low^ expression resulted in higher suppressive function in vitro. **(A)** Gene and protein expression of *TNFRSF1B*/CD120b and *ITGB1*/CD29 on UMAP projection of non MAIT CD8^+^CD45RC^low/-^ T cells. Gene expressions are scaled from grey to blue for genes or from grey to green for proteins. Each dot corresponds to a single cell. **(B) Left panel:** Gating strategy to identify CD45RC^low/-^ and ^high^ cells among CD8^+^ T cells and subsets within CD8^+^CD45RC^low/-^ or ^high^ T cells according to TNFR2 and CD29 expression by flow cytometry. **Right panel:** Frequency of the four subsets identified by TNFR2 and CD29 expression in CD8^+^CD45RC^low/-^ and ^high^ T cells. Statistical differences between groups were calculated using Mann-Whitney two-tailed test. *p value >0.05. **(C)** Normalized Mean Fluorescence Intensity (MFI) of TNFR2 and CD29 expression in CD8^+^CD45RC^low/-^ FOXP3^+^ or FOXP3^-^ T cell subsets normalized to isotype control. **(D)** Bar plot representing the percentage of proliferation of responder T cells in presence of subset of CD8^+^ Tregs at ratio Teff:Treg 1:1 normalized to the percentage of proliferation in absence of Tregs. Results are expressed as mean ± SEM (n= 6). Statistical differences between groups were calculated using Mann-Whitney two-tailed test. *p value > 0.05.

## DISCUSSION

In transplantation and inflammatory pathologies, one present major challenge is to develop new strategies to control inflammation with more specificity and less toxicity compared to immunosuppressive standard treatments. Recent clinical trials using CD4^+^ Tregs as cell therapy demonstrated absence of toxicity and hints of efficacy ^32^. CD8^+^ Tregs present at least equivalent suppressive activity compared to their CD4 counterparts ^7^. Notably in transplantation, CD8^+^ Tregs can take advantages of direct antigen presentation by MHC-I graft cells over the time as well as to donor APCs. In contrast, CD4^+^ Tregs can only become activated through direct presentation by donor APCs and transient activation of endothelial cells by MHC-II presentation. Expanded CD8^+^ Tregs (CD3^+^CD56^-^CD8^+^CD45RC^low/-^ Tregs) will be for the first time used as cell therapy for solid organ transplantation in the Eight-Treg clinical trial within the Reshape consortium (http://www.reshape.org). Investigations are ongoing to improve the therapeutic properties of CD8^+^ Tregs, including the use of CARs ^14^ and genetic cell modifications to improve persistence and stability ^33^. These efforts benefit from deeper characterization and a better understanding of CD8^+^ Tregs that is now possible with the development of single-cell analysis and new bioinformatic tools. In addition, understanding key markers of CD8^+^ Tregs will help monitor outcome of patients after transplant or with autoimmune diseases to adapt their treatment, and predict the efficacy of CD8^+^ Treg therapy in the upcoming phase I/IIa trial as part of the ReSHAPE consortium ^33^.

To the best of our knowledge, here we reported the first single cell RNA-seq dataset of human CD8^+^ Tregs. The data presented here characterize the transcriptomic heterogeneity inside CD8^+^ T lymphocytes from peripheral blood with the confirmation of 2 distinct subsets of cells: CD8^+^CD45RC^high^ and CD8^+^CD45RC^low/-^ T cells, highlighting CD45RC as a marker of pro-inflammatory vs pro-regulatory populations, as shown previously by us and by others ^9–11,13^. Surprisingly, among markers reported to identify CD8^+^ Tregs, only CD45RC was specific of one cluster. While TNFR2 and CD28 profiles may correlate, FOXP3, CD122, CD25… did not. This suggests the co-existence of several subsets of Tregs and explain the lack of consensus regarding their identity. This delineation of pro-inflammatory vs pro-regulatory populations is critical for the generation of a new therapeutic strategy as we have done with anti-CD45RC monoclonal antibodies in preclinical models of organ Tx ^9^, aGVHD ^34^, and APECED ^10^. Using the data generated here, pro-inflammatory or pro-regulatory specific targets could be identified for autoimmune diseases or cancer. A previous comparison between transcriptomes of CD8^+^CD45RC^high^ and CD8^+^CD45RC^low/-^ T cells was already done by 3’DGE-seq in both human and rat ^7,9^. This is the first time that their transcriptomic differences were analyzed using single cell RNA-seq data coupled to CITE-seq and TCR analysis. We were thus able not only to identify of new membrane markers, but also transcription factors. It has been clearly established that CD8^+^ T cells with regulatory properties are within the CD8^+^CD45RC^low/-^ T cells fraction ^6,7,9^. We excluded cells with more innate properties such as NKT cells excluded on CD56^+^CD16^+^ protein expression by CITEseq and MAIT cells based on CD161^+^ cells and TRAV1-2^+^ TRAJ33^+^ expression, to further focus on the CD8^+^CD45RC^low/-^ T cells pool, without losing any chance to detect CD56^+^ Tregs ^35,36^ or CD161^+^ Tregs ^37^. Thus, with this dataset, a new analysis could be conducted on these cell populations, but is out of the scope of this manuscript.

Interestingly, this bioinformatic work highlighted three interesting cell clusters within CD8^+^CD45RC^low/-^ T cells and in cell cluster 2, we observed higher levels of several genes associated with Tregs such as TNFR2 *(TNFRSF1B)*, PD1 *(PDCD1)*, *FASLG*, *IFNG*, and *FGL2* and lower levels of genes associated with T effector cells (*CD28*, *CD27* and *CD127*). We selected two genes encoding for membrane proteins with previous occurrence of association with Tregs, notably TNFR2 ^31,38–40^ and CD29 ^41^ to further specifically isolate this cell cluster for functional studies. We identified four subsets based on the expression of these two markers within peripheral blood fresh CD8^+^CD45RC^low/-^ T cells and demonstrated that the CD8^+^CD45RC^low/-^TNFR2^+^CD29^low^ T cell subset corresponded to the most suppressive CD8^+^ Tregs. Of note, previously identified markers like GITR tested to discriminate subsets within CD8^+^CD45RC^low/-^ Tregs required overnight polyclonal stimulation to achieve a similar 80% suppression at ratio Teff:Treg 1:1 ^7^. In contrast, here we show that isolation on TNFR2^+^CD29^low^ expression could achieve 90% suppression at similar ratio at steady state (without pre-stimulation). This association of suppressive function with TNFR2 expression is of great interest since several agonists targeting TNFR2 are under development and could lead to improve CD8^+^ Treg functional properties ^42,43^. We have shown that blocking TNFR2 signaling partially inhibited suppression mediated by CD8^+^ Tregs but we also previously showed that cytokines were also involved. We also performed experiments with a TNFR2 agonist to understand the role of TNFR2 for CD8^+^ Treg function, although the effect was modest (around 20 %), it correlates with the expression of TNFR2 within total CD8^+^CD45RC^low/-^ T cells. We observed that the CD8^+^CD45RC^low/-^TNFR2^+^ subset is CD28^-^ correlating with work previously described by others ^44^. Our team demonstrated that expression of IFN-γ and IL-10 is important for CD8^+^CD45RC^low/-^ Treg mediated suppression. Although IL-10 is not detected in this dataset, gene encoding for *IFNG* correlated with the cell cluster of interest.

Although *FOXP3* mRNA expression was not detected in enough cells of our dataset to draw conclusions, we observed that CD8^+^CD45RC^low/-^TNFR2^+^ cells expressed FOXP3 in higher amounts at the protein level compared to CD8^+^CD45RC^low/-^TNFR2^-^ cells. *IKZF2* (HELIOS) was in contrast detected and correlated with cell clusters 8 and 9. Interestingly, further transcriptomic analysis of these cell clusters demonstrated an interesting gene signature with cell cluster 9 resembling the TCR-biased CD8^+^ KIR2DL3^+^ Treg cell subset described by the recent work of the Davis’group ^4^ indicative of a CD8^+^ Treg subset involved to suppress pathogenic T cells in infectious diseases and in particular SAR-CoV-2 based on the TCR analysis. The overall TCR analysis of the other clusters revealed no specific public clonal TCR bias and no differences between clusters raising the question of their thymic origin.

Even if CD8^+^CD45RC^low/-^TNFR2^+^CD29^low^ cells represent a small population that could be difficult to isolate and amplify for cell therapy, they could serve as biomarker or biotarget and be expanded *in vivo* using agonists antibodies for autoimmune diseases or in the context of transplantation or using inhibitors to deplete or inhibit their function in cancer or infectious diseases.

CD29 is widely expressed in human body such as epithelial cells, has been reported for a role for attachment of cells to matrix and signal transduction, but its role in T cell function is not clear ^45,46^. CD29 has been identified to correlates with cytotoxic potential both in CD8^+^ T cells ^47^ and CD4^+^ T cells ^48^. Thus, its relevance and role of Tregs should be further investigated.

Finally, additional membrane markers were identified, such as *IL6R*/*IL6ST* (CD126/CD130), and should be investigated to better comprehend and define CD8^+^ Treg subsets that could have higher stability in inflamed environment. As for CD4^+^ Tregs for which studies on heterogeneity revealed distinct subsets ^49^ with different functions playing a particular role in different pathologic situations ^50^, it is highly probable that there are several populations of CD8^+^ Tregs with specific properties and playing a role in different situations. Understanding this heterogeneity could lead to specific therapeutic approaches in specific human conditions. Therefore, profiling CD8^+^ T cells in patients suffering from AID or transplanted versus patients suffering from cancer could help understand the importance/role in pathology of the Treg subsets identified in HV and inform on their stability (disappearance or clonal expansion?).

To conclude, using scRNAseq and bioinformatic methodology, we portrayed the heterogeneity of CD8^+^CD45RC^low/-^ Tregs and identified among thousand genes 2 genes encoding for membrane markers that are usable as biomarkers and promising targets for optimizing CD8^+^ Treg cell therapy in transplantation and autoimmune diseases.

## MATERIALS AND METHODS

### Isolation of human peripheral blood mononuclear cells (PBMC) of healthy volunteers

Blood samples from healthy volunteers were collected in EthyleneDiamineTetraAcetic acid (EDTA) coated tubes for omic analyses or from buffy coats at Etablissement Francais du Sang (EFS) for suppression assays. PBMC were isolated with Ficoll-Paque density gradient centrifugation (Eurobio, Courtaboeuf, France). Red blood cells and platelets were removed with hypotonic lysis solution and centrifugation.

### Staining and cell sorting of cells

PBMC were diluted in Phosphate Buffered Saline (PBS) – FCS 2% -EDTA 2 mM and labelled with surface antibodies. anti-CD3 (BD Biosciences, SK7 clone), anti-CD4 (BD Biosciences, RPA-T4 clone), anti-CD45RC (IqProducts, MT2 clone) and with 4′,6-diamidino-2-phenylindole (DAPI). For suppression assay, anti-CD25 (BD Biosciences, M-A251clone) mAbs was added. Lymphocytes were selected according to their morphology in FSC-A and SSC-A, doublets cells were excluded and DAPI^+^ cells were removed. DAPI^-^CD3^+^CD4^-^ were sorted for CITE seq, DAPI^-^CD3^+^CD4^-^CD45RC^low/-^ T cells and DAPI^-^CD3^+^CD4^+^CD25^-^ T cells were sorted for suppression assay. In some cases, four subsets in CD8^+^CD45RC^low/-^ Tregs were labelled and sorted according to the expression of TNFR2^+/-^ (Miltenyi Biotec, REA520) and CD29^low/high^ (Miltenyi Biotec, REA1060). Purity was greater than 95%.

Cells were sorted with FACS ARIA II cell sorter (BD Biosciences) and data were analyzed with FLOWJO software (Tree Star, Inc., Ashland, OR, USA).

### Single cell RNA-sequencing of blood CD8^+^ T cells

Freshly sorted CD8^+^ T lymphocytes of four healthy volunteers were labelled with anti-human Hashtag antibodies TotalSeq-C (Biolegend, San Diego, CA) and were pooled together. Cell viability was > 90 %. Then, cells were labelled with 30 CITE-seq monoclonal antibodies including antibody against CD45RC (ABIS clone, homemade), CD120b (3G7A02 clone), HLADR (L243 clone), CD39 (A1 clone), CD357 (108-17 clone), CD279 (EH12.2H7 clone), CD152 (BNI3 clone), TIGIT (A15153G clone), CD223 (C9B7W clone), CD70 (113-16 clone), CD137 (4B4-1 clone), CD226 (11A8 clone), CD25 (BC96 clone), CD71 (CY1G4 clone), CD134 (Ber-ACT35 (ACT35) clone), CD11c (S-HCL-3 clone), CD16 (3G8 clone), CD56 (5.1H11 clone), CD28 (CD28.2 clone), CD45RA (HI100 clone), CD45RO (UCHL1 clone), CD103 (Ber-ACT8 clone), CD178 (NOK-1 clone), CD29 (TS2/16 clone), CD122 (TU27 clone), CD127 (A019D5 clone), CD126 (UV4 clone), CD130 (2E1B02 clone), CD21 (Bu32 clone) and CD197 (G043H7 clone). Single cell libraries were performed according to Chromium Next GEM Single Cell 5’ Reagent Kits v2 protocol (10X genomics, San Francisco, CA). Cell suspension were loaded onto chromium single cell chip K in 4 different well (20 000 cells per well) and run immediately on the Chromium controller (10X genomics, San Francisco, CA). Three libraries were prepared: one for mRNA (RNA), one for barcoded antibodies (HTO and CITE-seq) and one for TCR (VDJ-seq). They were sequenced with NovaSeq 6000 (Illumina, San Diego, CA) according to 10X genomics recommendations.

### Single cell RNA-sequencing data analysis

#### 1/ primary analysis

FASTQ files, first generated from BaseCalling (BCL) files with CellRanger package (6.1.2), were demultiplexed and aligned to human reference genome (hg38). CITE-seq-Count function (1.3.4) was used to count antibody hashtag sequences.

#### 2/ secondary analysis

Count matrixes were analyzed with Seurat R package (4.1.1). Cells with more than 10% of mitochondrial genes were excluded. Gene expression was log normalized and scaled. HTO expression was normalized and demultiplexed. Doublet cells and negative cells for HTO were removed. Downstream analysis was performed on 15718 single cells: 7535 fresh CD8^+^ T lymphocytes. On average 1454 genes were expressed per cell in fresh CD8^+^ T cells. A nonlinear dimensionality reduction Uniform Manifold Approximation and Projection (UMAP) and a clustering were performed to visualize distinct cell clusters with a resolution of 1 for total CD8^+^ T cells, 0.8 after MAIT cells exclusion, and 0.5 for CD8^+^CD45RC^low/-^ T cells only. Differentially expressed genes were calculated using “wilcox” method to characterize marker genes of each cell cluster implemented in the Seurat package’s “FindAllMarkers” or “FindMarkers” functions. CITE-seq antibodies gates were set up according to expression profiles detected by flow cytometry with negative and positive fractions and for some CITE-seq antibodies including anit-CD45RC with low and high fractions.

#### 3/ TCR repertoire analysis

This dataset contained matched TCR for each single cell. For TCR analysis, scRepertoire package was used (1.7.2). TCR repertoire was analyzed by studying the length of the amino acid sequence of the CDR3 regions of TRA and TRB chains and by exploring clonotypes expansion. The interconnectivity between clusters regarding TCR sequences/clonotypes sequences were analyzed and visualized using chord plots. Finally, the iterations of each TCR sequence/clonotype in each cell cluster is presented based on the absolute frequency.

### Suppressive assays

CD8^+^ Treg cell subsets sorted according to TNFR2 and CD29 expression were cultured in RPMI 1640 medium supplemented with glutaMAX, 1% NEAA, 1% sodium pyruvate, 1% Hepes 1% Penicilin/Streptomycin, and 5% human AB serum with autologous CD4^+^CD25^-^ conventional/responder T cells labelled with CFSE and irradiated allogenic antigenic presenting cells (APC). When indicated, blocking or anti-TNFR2 monoclonal antibody (358408, Biolegend) or its isotype control (400544, Biolegend) or the human TNFR2-specific agonist TNC-scTNF(143N/145R), ^30^ were added. After 5 days of co-culture at 37°C, 5% CO_2_, CFSE fluorescence intensity was measured in CD3^+^CD4^+^DAPI^-^ T cells with BD FACS CANTO II (BD Biosciences) to measure the proliferation of CD4^+^CD25^-^ effector T cells with or without the presence of CD8^+^CD45RC^low/-^TNFR2^+^ Tregs or total CD8^+^CD45RC^low/-^ Tregs.

### Statistics

Non-parametric Mann Whitney test was performed to compare TNFR2 and CD29 MFI in FOXP3^+^ and FOXP3^-^ CD8^+^CD45RC^low/-^ T cells. Two-way ANOVA test was performed to compare TRA and TRB chain frequency in TNFR2^+^ and TNFR2^-^ CD8^+^CD45RC^low/-^ T cells. Volcano plot showing – log10 p-value and log2 fold change in y and x axis respectively. P-values were calculated with FindMarkers function of Seurat R package. Mann Whitney test, non-parametric, p value two tailed was performed to compare CD8^+^ Treg subsets suppressive functions at ratio Teff:Treg 1:1. Two-way row-matched ANOVA was performed to compare suppressive function of CD8^+^CD45RC^low/-^TNFR2^+^ T cells with blocking antibody anti-TNFR2 or isotype control and with STAR2 or isotype control in a range of cell ratio.

## Acknowledgments

We thank Sonia Salle for blood sampling. This work was partially funded by the Labex IGO program supported by the National Research Agency via the investment of the future program ANR-11-LABX-0016-01. This work was supported by an “Etoiles Montantes” from Pays de la Loire to C.G and the Deutsche Forschungsgemeinschaft (DFG, German Research Foundation) – project number 324392634 – TRR221 project B02 to H.W. This work was also realized in the context of the support provided by the Fondation Progreffe. This project has received funding from the European Union’s Horizon 2020 research and innovation program under grant agreement No 825392 “RESHAPE”.

## FIGURE LEGENDS

**Suppl. Figure 1:**
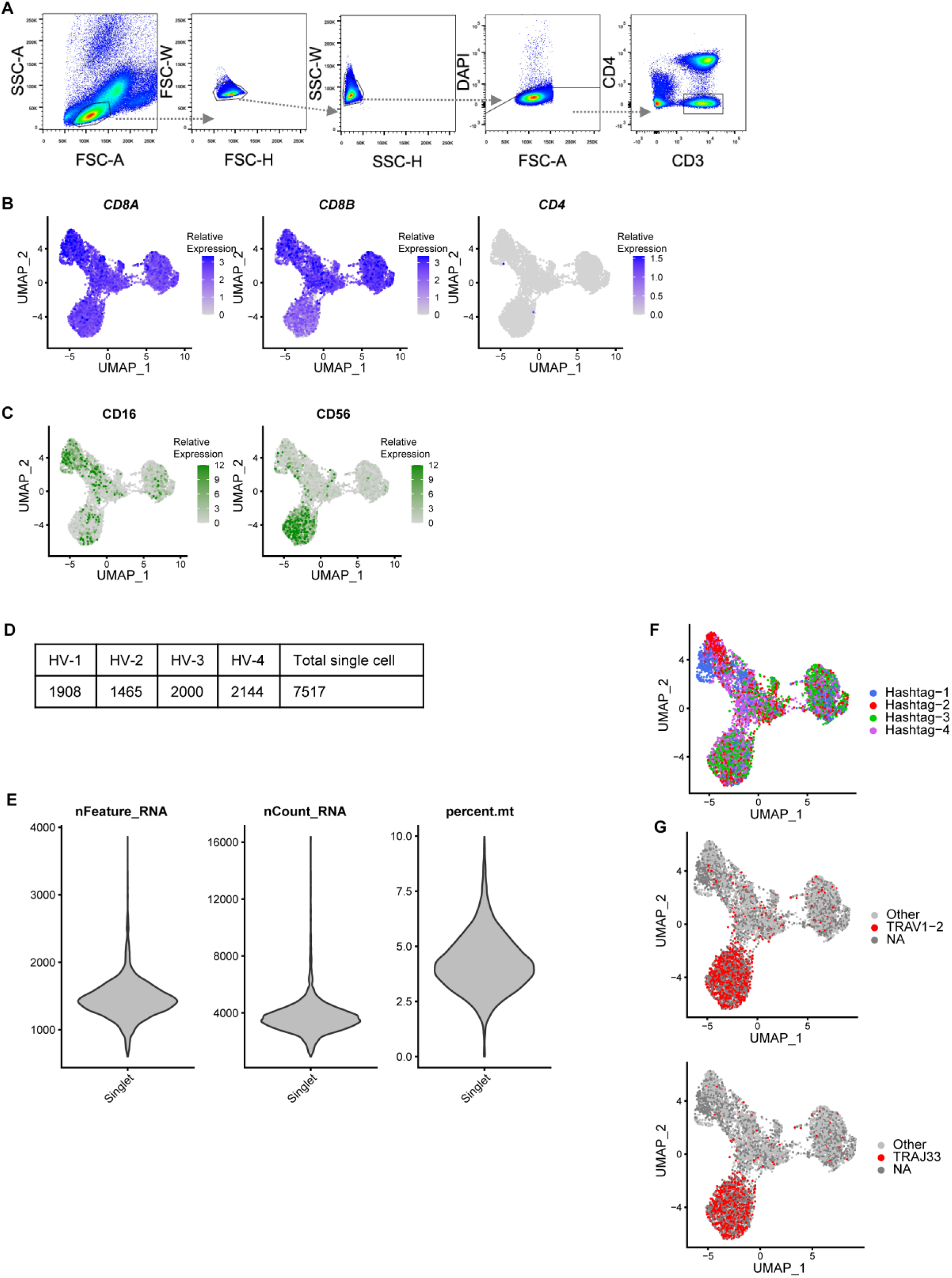
**(A)** Cell sorting strategy of CD8^+^ T cells. CD8^+^ T cells were defined by gating on lymphocytes morphology, FSC and SSC singlets, living cells, CD3^+^CD4^-^. **(B)** Feature plots showing gene expression of *CD8A*, *CD8B* and *CD4*. Gene expressions are scaled from grey to blue. Each dot corresponds to a single cell. **(C)** Feature plots showing protein expression of CD16 and CD56. Gene expressions are scaled from grey to green. Each dot corresponds to a single cell. **(D)** Table of the number of single cells per healthy volunteer (HV). **(E)** Violin plot representing the number of genes detected in each single cell, the number of RNA molecules sequenced in each single cell and the percentage of mitochondrial genes in each single cell for the all 4 HV. **(F)** UMAP showing the affiliation of the cells to each of the healthy volunteers. **(G)** TCR sequences of TRAV1-2 and TRAJ33. Cells expressing these chains of the TCR are in red, cells in light grey expressing other chains and for cells in dark grey no TCR chain were detected. **(H)** Top 10 more represented TCR sequences among the total CD8^+^ single cell dataset.

**Suppl. Figure 2:**
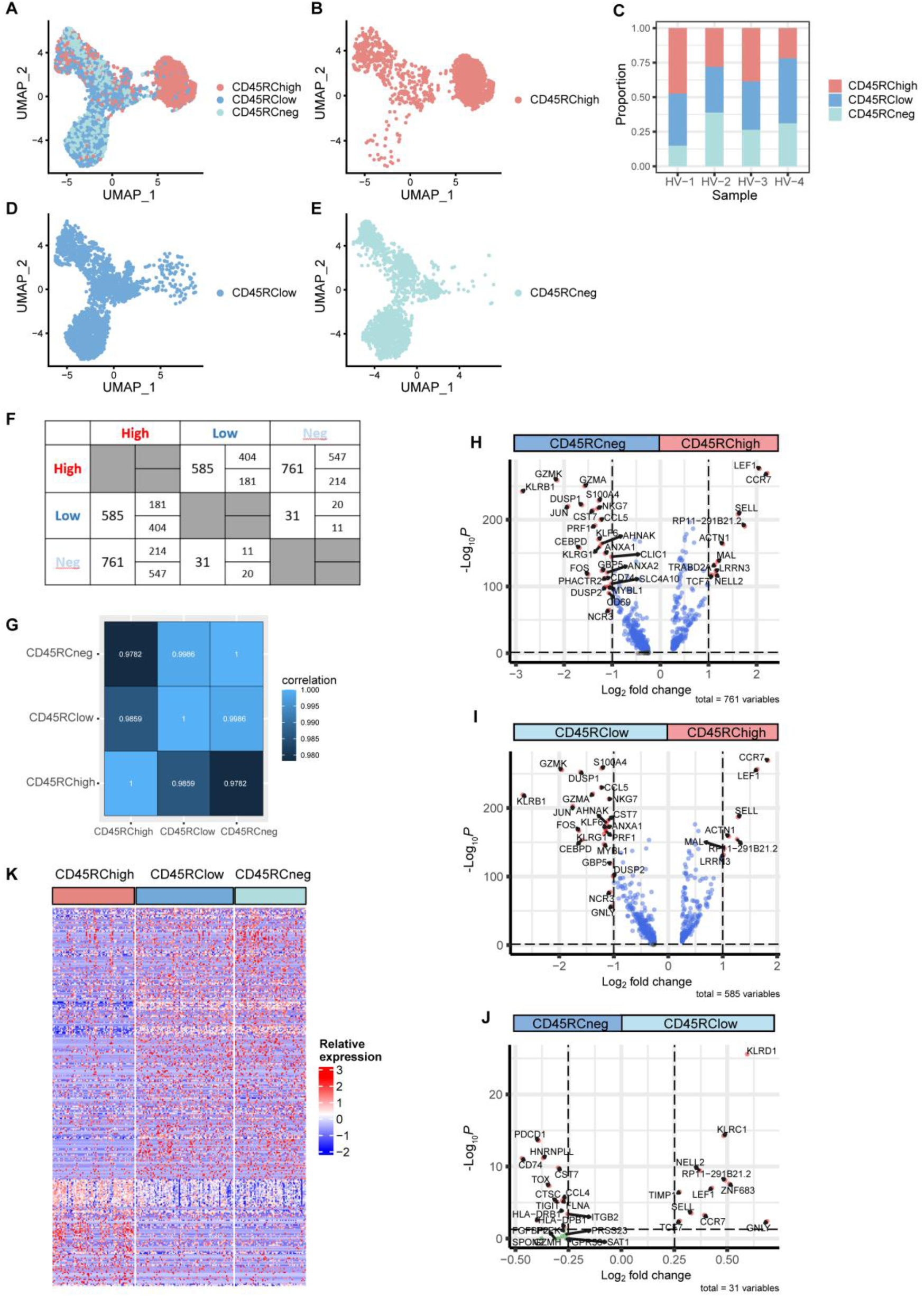
**(A)** Definition of CD8^+^CD45RC^high^ (orange), ^low^ (dark blue) and ^neg^ (light blue) cell subsets with anti-CD45RC CITE-seq antibody on the UMAP plot. Each dot corresponds to a single cell. **(B-D)** CD8^+^CD45RC^high^ cells (B), CD8^+^CD45RC^low^ cells (C) or CD8^+^CD45RC^neg^ cells (D) projection on umap plot (A). **(E)** Proportion of CD8^+^CD45RC^high^ (orange), ^low^ (dark blue) and ^neg^ (light blue) cell subsets in each healthy volunteer. **(F)** Table of the number of differential expressed genes between CD8^+^CD45RC^high^, ^low^ and ^neg^ cell subsets. **(G)** Correlation heatmap reporting correlation coefficient between CD8^+^CD45RC^high^, ^low^ and ^neg^ cell subsets. The lighter the blue is, the greater the correlation is. **(H-J)** Volcano plots showing differentially expressed genes between cell cluster CD8^+^CD45RC^neg^ and CD8^+^CD45RC^high^ T cells (H) or CD8^+^CD45RC^low^ and CD8^+^CD45RC^high^ T cells (I) or CD8^+^CD45RC^neg^ and CD8^+^CD45RC^low^ T cells (J). The thresholds have been set to p-value adjusted <0.05 and fold change >1 (H-I) or >0.25 (J). Red dot corresponds to genes that have exceeded both threshold, and blue dot correspond to genes that have a significant p-value adjusted but not fold change. The top most differentially expressed genes were labelled. **(K)** Heatmap of genes encoding for differential expressed genes between CD8^+^CD45RC^high^, ^low^ and ^neg^ cell subsets. Columns correspond to cell clusters and rows correspond to genes. Expression values were scaled per gene. Blue color represents less expressed genes and red color represents more expressed genes.

**Suppl. Figure 3:**
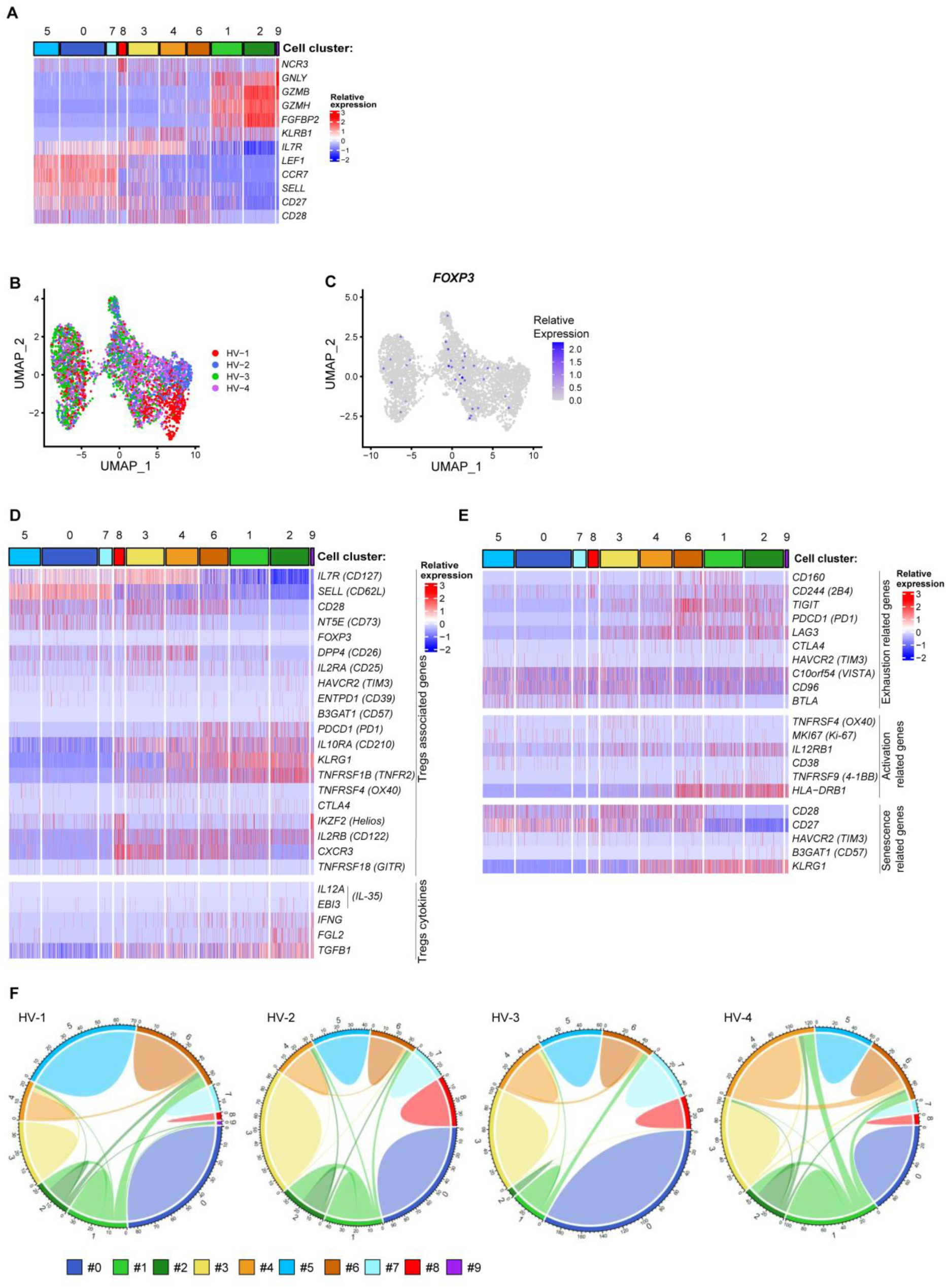
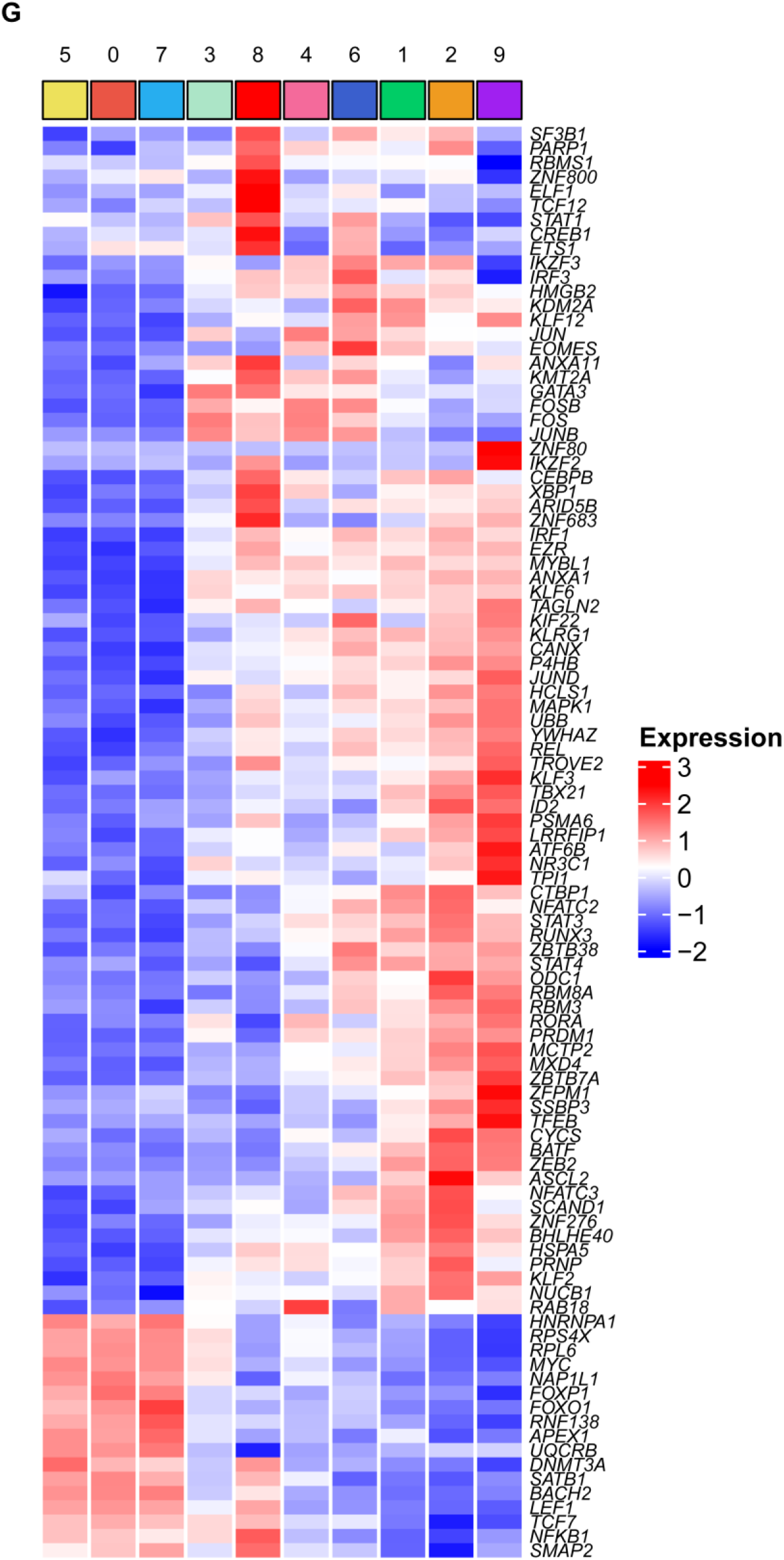
**(A)** Heatmap of selected genes for defining clusters identity. Columns correspond to cell clusters and rows correspond to genes. Expression values were scaled per gene. Blue color represents low expressed genes and red color represents highly expressed genes. **(B)** UMAP showing the affiliation of the non MAIT CD8^+^ T cells to each of the healthy volunteers. **(C)** Feature plot showing gene expression of *FOXP3* in non MAIT CD8^+^ T cells. Gene expression is scaled from grey to blue. Each dot corresponds to a single cell. **(D)** Heatmap of genes encoding for membrane, transcription factor or cytokines associated with Tregs. Columns correspond to cell clusters and rows correspond to genes. Expression values were scaled per gene. Blue color represents less expressed genes and red color represents more expressed genes. **(E)** Heatmap of genes encoding for genes related to exhaustion, activation or senescence. Columns correspond to cell clusters and rows correspond to genes. Expression values were scaled per gene. Blue color represents less expressed genes and red color represents more expressed genes. **(F)** Chord plot showing TCR clonotypes sharing between cell clusters for each healthy volunteer. Numbers from 0 to 9 and colors indicate the cell cluster defined in Fig 2A. **(H)** Heatmap showing the differentially expressed genes between cell clusters encoding for transcription factors min.pct = 0.25, logfc.threshold = 0.2). Columns correspond to cell clusters and rows correspond to genes. Expression values were averaged per cell cluster and scaled per gene. Blue color represents less expressed genes and red color represents more expressed genes.

**Suppl. Figure 4:**
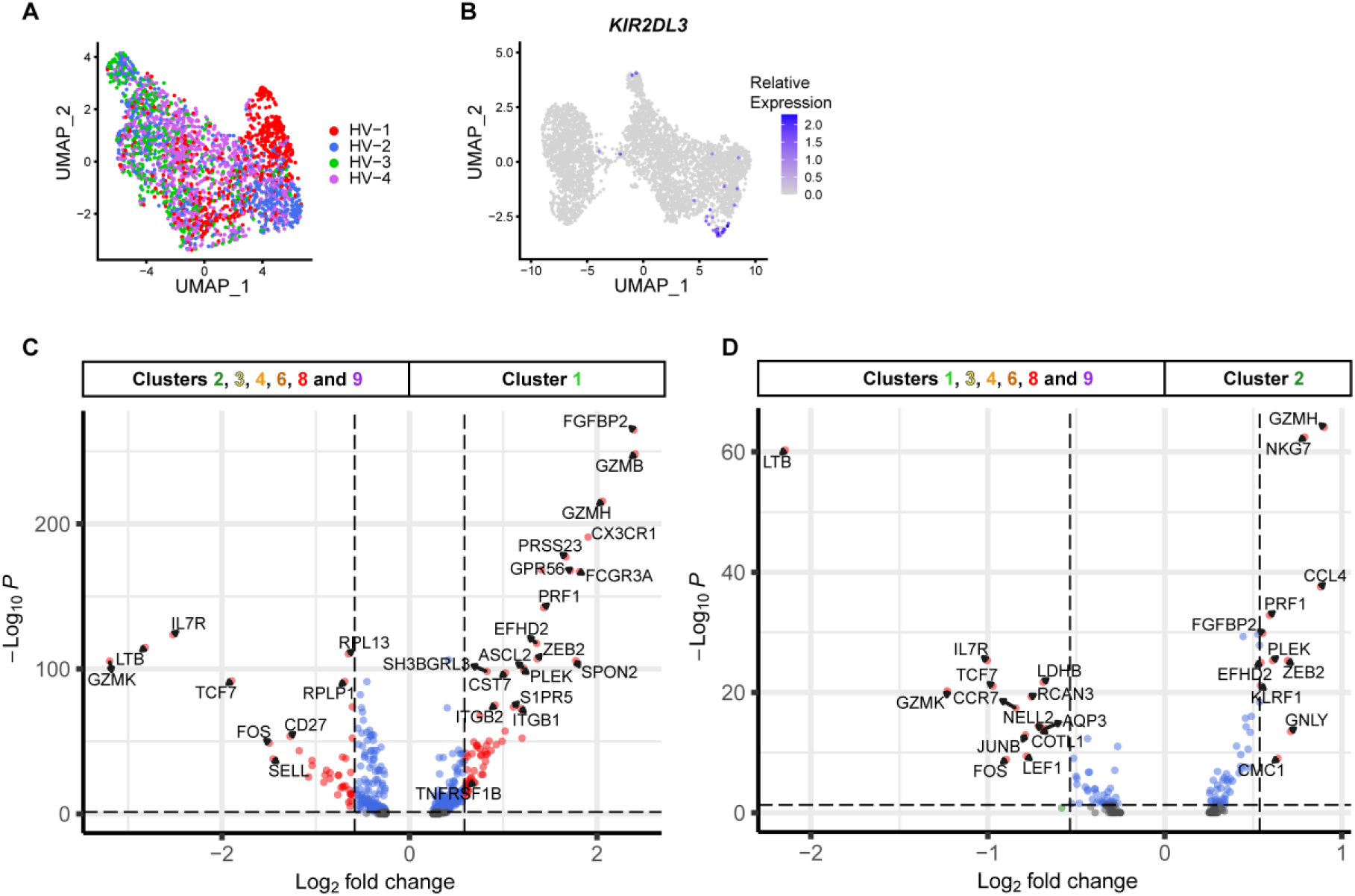
**(A)** UMAP showing the affiliation of the non MAIT CD8^+^CD45RC^low/-^ T cells to each of the healthy volunteers. **(B)** Feature plot showing gene expression of *KIR2DL3* in non MAIT CD8^+^T cells. Gene expression is scaled from grey to blue. Each dot corresponds to a single cell. **(C-D)** Volcano plot showing differentially expressed genes between cell cluster 1 vs all other cell clusters **(C)**, between cell cluster 2 vs all other cell clusters **(D)**. The thresholds have been set to p-value adjusted <0.05 and fold change >1.5 **(C)** or 1.45 **(D)**. Red dot corresponds to genes that have exceeded both threshold, and blue dot correspond to genes that have a significant p-value adjusted but not fold change. *TNFRSF1B* gene **(C)** and the top most differentially expressed genes were labelled **(C-D)**.

**Suppl. Figure 5:**
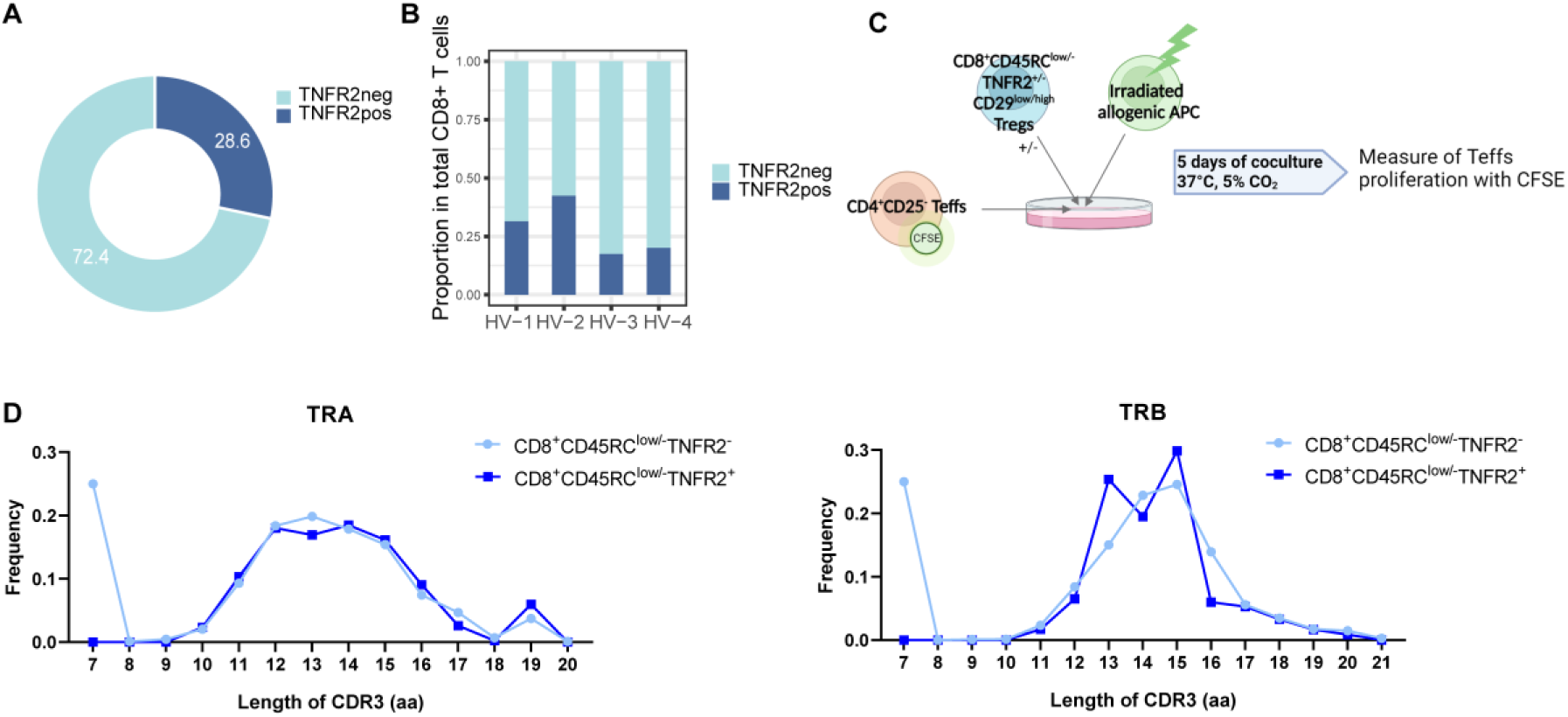
**(A)** Percentage of TNFR2^+^ and TNFR2^-^ CD8^+^CD45RC^low/-^ T cells represented as a donut chart. TNFR2^+^ CD8^+^CD45RC^low/-^ T cells are represented in dark blue and TNFR2^-^ CD8^+^CD45RC^low/-^ T cells are represented in light blue. **(B)** Proportion of TNFR2^+^ and TNFR2^-^ CD8^+^CD45RC^low/-^ T cells in each healthy volunteer. TNFR2^+^ CD8^+^CD45RC^low/-^ T cells are represented in dark blue and TNFR2^-^ CD8^+^CD45RC^low/-^ T cells are represented in light blue. **(C)** Suppressive assays workflow. Proliferation of responder T cells (CD4+CD25- T cells) labelled with CFSE in presence of irradiated allogenic Antigen Presentating Cells (APC) with or without subset of CD8^+^ Tregs is measured 5 days after co-culture at 37°C with 5% of CO_2_. **(D)** Length distribution of the CDR3 sequences in amino acids per cell cluster (aa) for TRA chains (left part) and for TRB chains (right part). CD8^+^CD45RC^low/-^TNFR2^-^ T cells are represented with light blue and CD8^+^CD45RC^low/-^TNFR2^+^ are represented in dark blue.

**Suppl. Table 1:**
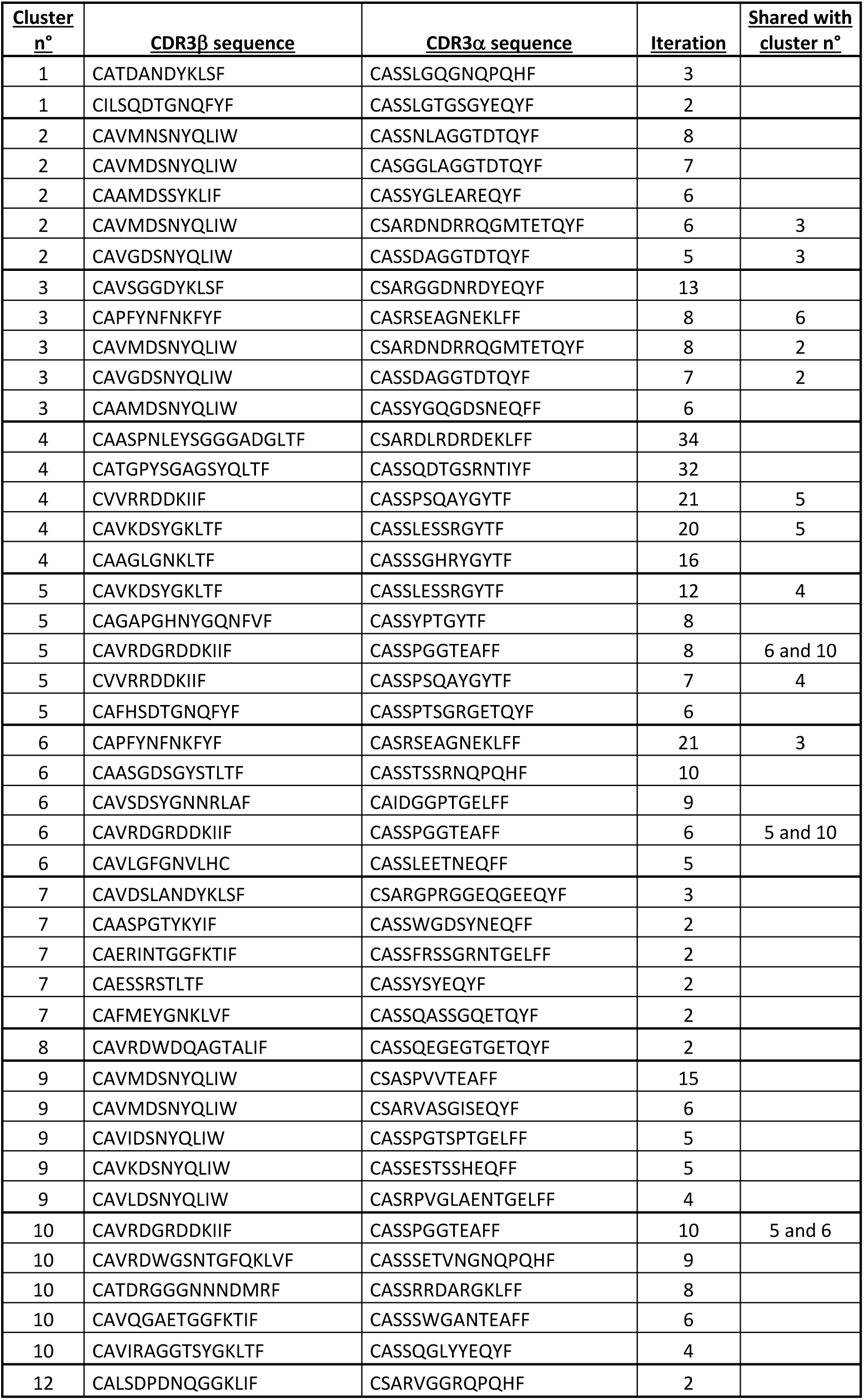
Most frequent CDR3β used sequences within clusters.

